# Synthetic-data augmented calibration for expert-informed rare disease models

**DOI:** 10.64898/2026.05.18.725833

**Authors:** Hanning Yang, Timo Rachel, Tim Litwin, Meropi Karakioulaki, Antonia Reimer-Taschenbrecker, Jens Timmer, Cristina Has, Harald Binder, Moritz Hess

## Abstract

Clinical data for rare diseases are sparse, noisy, and heterogeneous, complicating calibration of ordinary differential equation (ODE) models. Thus, we introduce a noise-robust calibration in latent space that combines expert-derived ODEs with learned latent representations. Our approach leverages synthetic ODE trajectories, augmenting our scarce observations to train a model-specific autoencoder representation and imputer. During calibration, observed and ODE-generated trajectories are compared in latent space, and ODE parameters are updated by minimizing their latent distance. In a controlled ABCDE simulation model, the imputer outperformed a carry-forward baseline for moderate parameter shifts, parameter recovery remained stable under random missingness, calibration remained robust to additional noise variables despite reduced downstream identifiability, and distinct dynamics formed visually separable latent trajectories. On a custom developed ODE model for real Epidermolysis Bullosa patients, the calibrated phenomenological model reproduced patient-level trajectories from sparse observations. Thus, we conclude that our latent-space calibration approach supports rare-disease modeling.

## Introduction

Clinical data in rare diseases such as Epidermolysis Bullosa (EB) are inherently complex [1], characterized by datasets with few patients with significant intra- and inter-patient variabilities of diagnostically relevant variables [2]. Even when patient data for rare diseases are available, their time-courses are typically only sparsely observed and some important clinical biomarkers might be missing entirely [2, 3]. Such inherent data scarcity renders purely data-based inference of patient disease courses or prediction treatment effects ineffective.

Much progress has been made in understanding diseases not only on a phenomenological, but also on a biologically more fundamental level. In systems medicine, biological processes are represented through molecular and multiscale models that integrate genes, proteins, pathways and clinical phenotypes [4]. Such approaches have been used, for example, to propose candidate mechanisms of Alzheimer’s disease through multi-omics network-interactions [5] or to predict treatment response to rheumatoid arthritis with a mechanistic differential equation model [6]. Even when not directly applicable to patient level data, the enhanced understanding of disease processes facilitates selection of clinically relevant biomarkers that can be included to build more appropriate phenomenological models.

Disease processes are often modeled by ordinary differential equations (ODE) because they align the possible dynamics of variables with our understanding of the process. Furthermore, ODEs often constitute the most natural way to describe the time-evolution of a system, because they describe the future behavior of the system given the currently observed state. This makes them a good fit for clinical settings, because the availability of prior patient history in terms of the relevant variables will often be limited.

The dynamical behavior implied by an ODE depends on the chosen values for the different model parameters. However, calibrating these model parameters from patient data in a rare disease setting runs into various difficulties. For one, it is not guaranteed that the chosen model structure represents the groundtruth; especially phenomenological models might contain clinical variables that are less important for the disease and thus mislead the calibration procedure. Another issue is the strong inter-patient heterogeneity that might drown out the signal in the intra-patient dynamics. Even once a model is calibrated, humans might struggle to visually infer important differences between patients from the trajectories of clinical variables due to a high number of variables involved.

While classical ODE modeling is especially powerful in mechanistic contexts and can also handle individual heterogeneity through mixed effects modeling [7], challenges with regards to phenomenological settings remain. A typical modeling process includes hypothesizing on the important variables and the structure of the model, assessing the fit of model predictions, and gathering data from carefully planned experiments to clarify the causal relationship between variables [8, 9]. Existing ODE modeling toolboxes such as **Data2Dynamics** [10], **dMod** [11], or **PyPESTO** [12] assume the model structure to be the ground truth model, which later can be reduced by model selection techniques such as AIC/BIC [13] data-based model reduction [14]. Such an approach is vulnerable to noisy and sparse data, because the parameter calibration can not properly distinguish the relevant from the irrelevant model variables, causing model and parameter identifiability issues [15, 16]. Given the nature of clinical data and the often missing causal knowledge in the rare disease context, phenomenological models are especially susceptible to uninformative variables.

Integrating concepts of artificial intelligence into existing ODE modeling approaches might help alleviate misspecifications of model structures. Model-discrepancy approaches quantify the mismatch caused by model calibration from the one caused by model misspecification [17], but might struggle with this separation with poor quality data. In a physics-informed context, the universal ODE approach [18] allows specifications of a core ODE that is augmented by neural network terms, which on the one hand enables manual specification of the important ODE links, but on the other hand comes with a flexibility that is difficult to properly constrain in the limited data setting. Other approaches utilize both autoencoders [19] and ODEs on the associated latent space [20, 21] to properly constrain the dynamic freedom of the model while alleviating the impact of noise, but stop short of utilizing an ODE in the observed space, on which experts can explicitly try to link relationships between variables.

We propose to combine both an initial guess ODE on the observed space as well as a latent space representation learned on likely synthetic patient trajectories to calibrate real patient trajectories in the presence of sparse patient data and a misspecified ODE (Figure 1). We make use of the initial guess ODE to generate synthetic data based on a distribution of initial conditions for the patient population. This dataset is subsequently used to train an autoencoder that learns patient-standardized correlations between clinical variables and also to train an imputation algorithm that leverages the knowledge in the ODE structure to better impute the missingness within the patient data. The ODE parameters are then calibrated to align the ODE trajectories with the real patient data based on their distance in latent space, which reduces the impact of noise variables and allows better discrimination of patient groups in latent space. We first demonstrate the performance of our approach on a simulated model with a known ODE structure and deliberate introduction of noise variables. We further apply our approach to real patient data for Epidermolysis Bullosa, where we accurately calibrate patient trajectories based on an expert-developed ODE.

**Figure 1:**
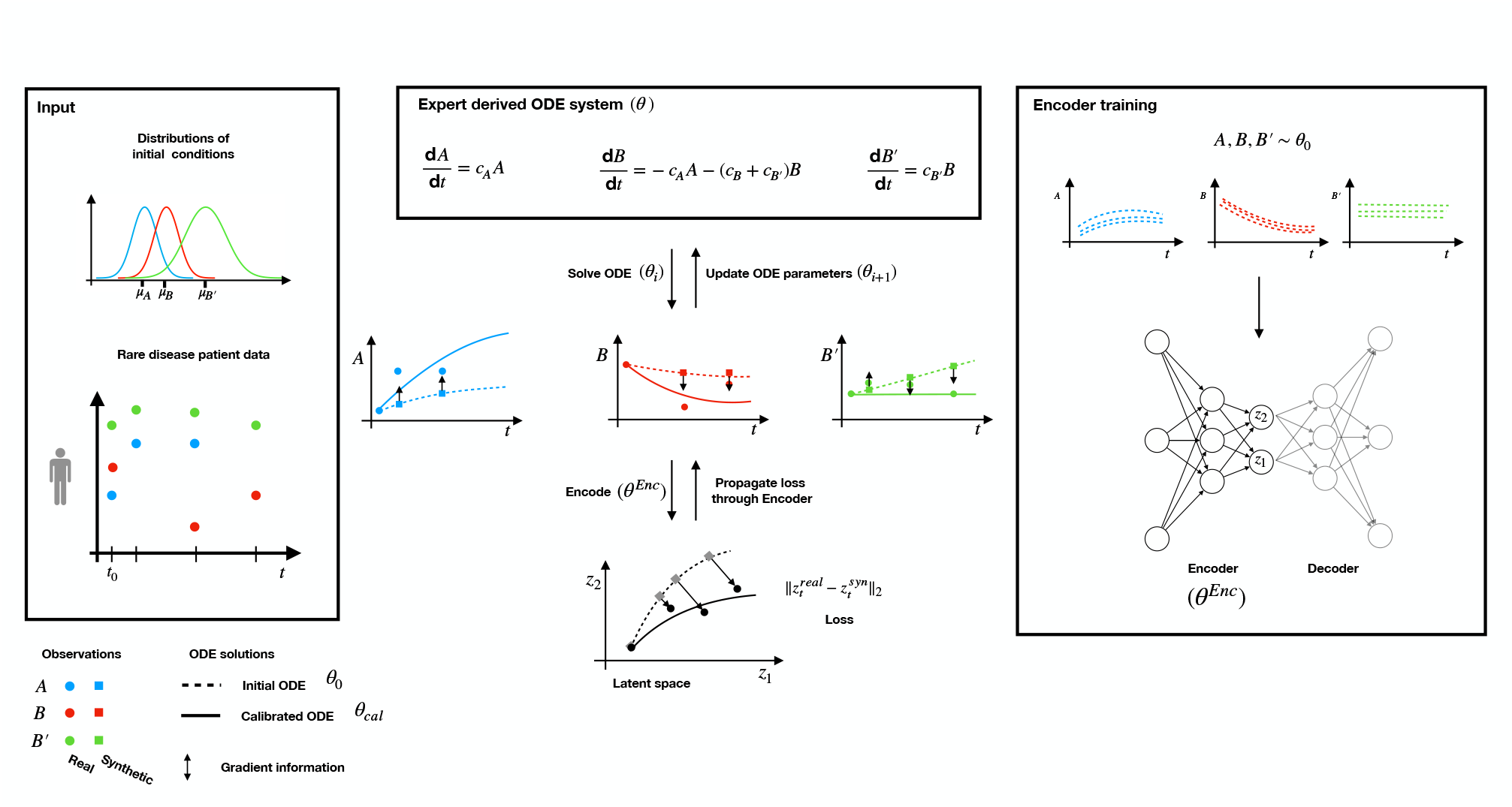
We calibrate ODEs from sparse observational patient data and patient-calibrated initial condition distributions. An initial ODE system with parameters *θ*_0_ is derived through expert input for the variables *A, B* and the misspecified noise variable *B*^′^. With this initial ODE system, a latent representation of the synthetic patient data is learned by an autoencoder. The ODE model parameters are iteratively updated by aligning the synthetic data generated with *θ*_*i*_ with the real data within the learned latent space. Gradients (arrows) computed for the loss are propagated back through the encoder weights and the ODE solver to update the ODE parameters in an iterative manner (*θ*_*i*_ → *θ*_*i*+1_).

## Results

### Calibration Remains Robust to Observation Noise and Missingness

As a first controlled evaluation of the calibration workflow, we used the simulated ABCDE model (Fig. 2(a)) (see Eq. (1) in Methods) with known ground-truth parameters. It is a simple linear compartmental chain model, to demonstrate performance in a setting in which the variable relationships change non-linearly.

**Figure 2:**
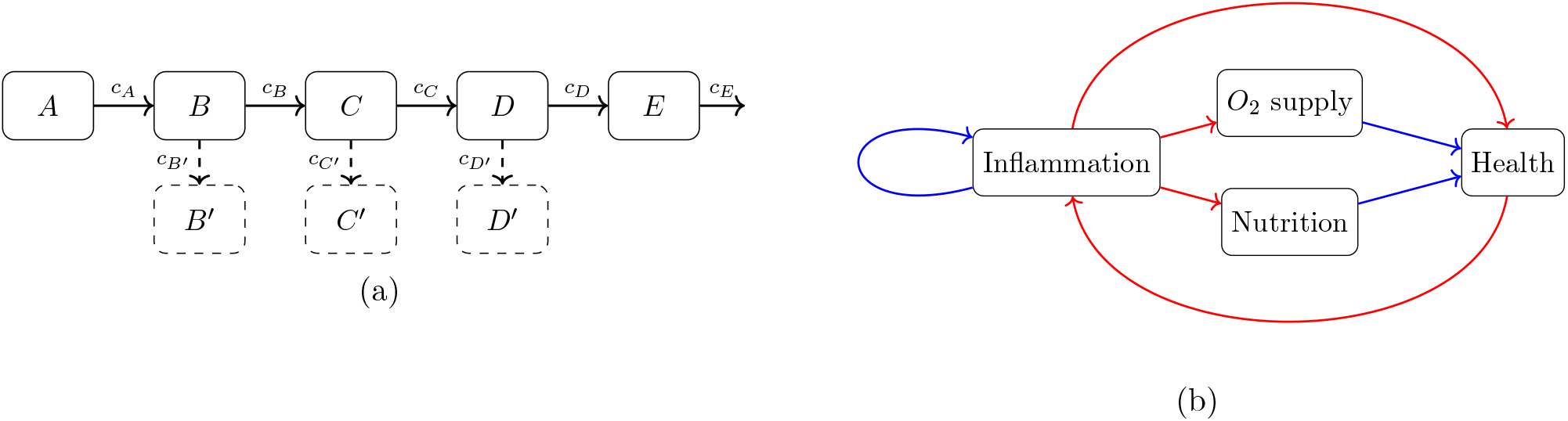
Visual representations of the **(a)** simulated ABCDE model and **(b)** Epidermolysis Bullosa model. The ABCDE model represents a chain of mass-conserving conversions, where the dashed arrow reflect alternative versions of the model with additional states for testing the importance of noise variables. The Epidermolysis Bullosa model shows a conceptual loop for 4 phenomenological states, where red arrows represent that one variable causes a relative decrease on another variable while blue allows represent a relative increase.

To check the performance of our model systematically and check for internal consistency of our approach, we defined different simulation settings. We initialized our simulations with true base parameter sets in which all conversion parameters were set to 1 and have simulated the ground truth data from this set with 10% noise and implemented different levels of missingness completely at random (MCAR). We further assessed the sensitivity of our calibration approach to the quality of the initial guess by either drawing initial guess parameters with distance [0, 0.2] or [0.2, 0.5] from the ground-truth parameters. For each calibration setting, calibration was repeated across 50 independent runs.

We compared the mismatch between imputed value and the ground truth for both our imputer and a last observation carried forward imputation (LOCF) as a baseline imputation algorithm. LOCF replaces each missing value is by its previously observed value for the same patient and biomarker. Missing values before the first observation were filled by backward filling from the first available later value.

As shown in Fig. 3(a), the ODE-informed imputation model produced lower reconstruction errors than LOCF for small parameter perturbations across all tested MCAR rates. For larger parameter perturbations, imputation became less accurate for both approaches, indicating that the benefit of the ODE-informed imputer depends on the training ODE being sufficiently close to the data-generating dynamics.

**Figure 3:**
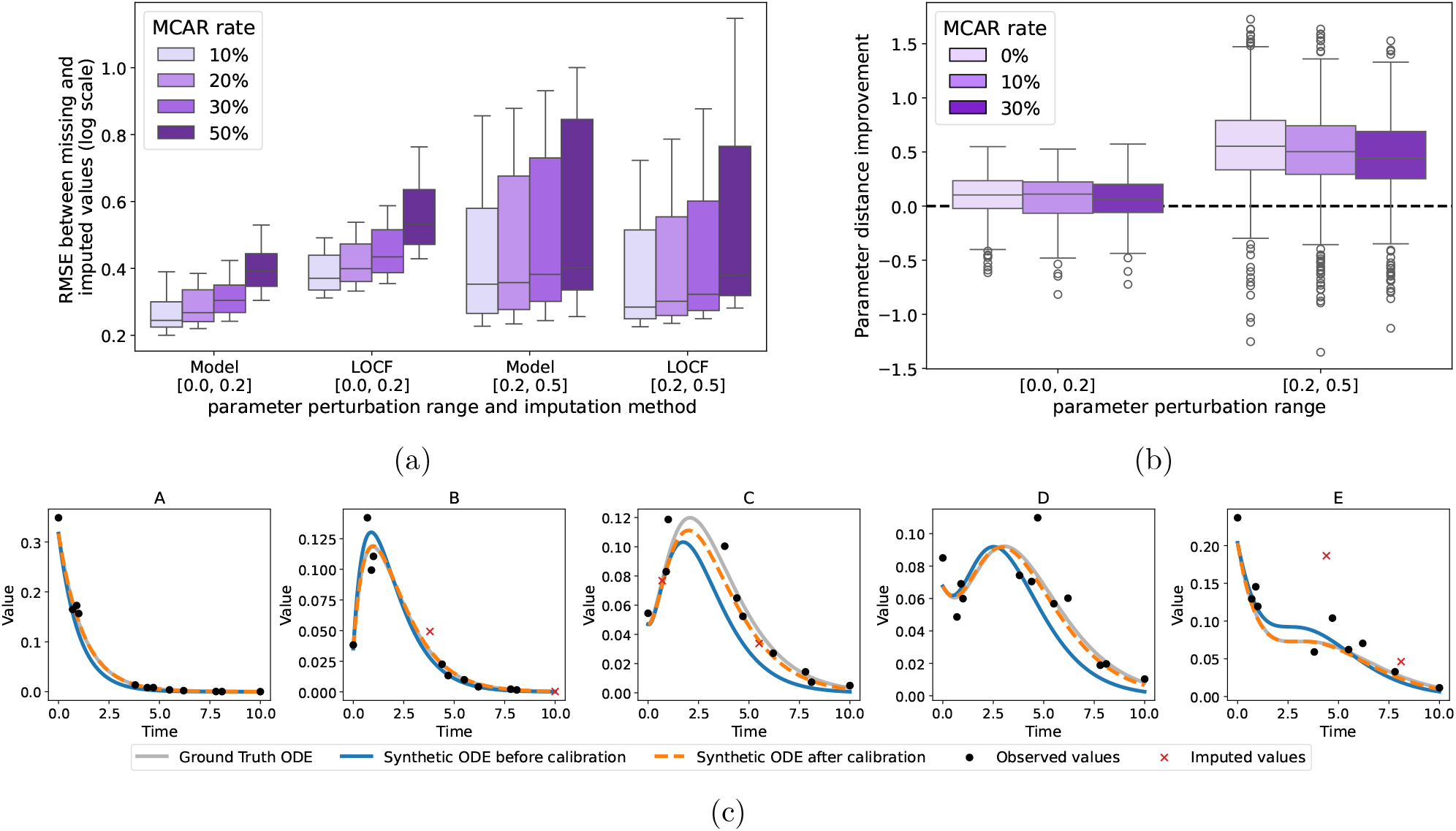
Performance of the calibration workflow on the simulated ABCDE model. **(a)** Imputation performance of the ODE-informed imputation model compared with last observation carried forward (LOCF) with backward filling for leading missing values. RMSE was computed between imputed and masked values on the log scale for different MCAR rates and two parameter pertubation ranges. **(b)** Parameter distance improvement after calibration. Positive values indicate that the calibrated core parameters *c*_*A*_–*c*_*E*_ are closer to the true values than the initial parameters. Boxplots compare complete data with 10% and 30% MCAR missingness for the two parameter pertubation ranges [0.0, 0.2] and [0.2, 0.5]. (C) Representative simulated patient trajectory showing observed values, imputed missing values, the ground truth ODE used for data generation, and the synthetic trajectories before and after calibration.

When imputation was included in the calibration workflow, recovery of the ground-truth parameters remained comparable between datasets without missingness and missingness up to 30% (Fig. 3(b)), which was observed for both small and larger parameter perturbations. The overall ability to retrieve the original ground-truth trajectories is illustrated in Fig. 3(c) for a representative patient trajectory for this simulation model.

Together, these results show that the workflow can handle observation noise and moderate MCAR missingness in the controlled ABCDE setting while retaining the ability to recover the ground-truth parameters and trajectories from flawed data.

### Calibration Ignores Noise Variables

To test whether latent-space calibration is robust to adding irrelevant clinical variables into the model, we extended the ABCDE model by adding three states that do not contribute to the original dynamics (Fig. 2). This lack of contribution is simulated by setting the conversion rates of these irrelevant states close to zero for the ground-truth data, such that the model does not know these states are irrelevant but has to infer this fact from the simulated data. We compared the base model with extended versions containing one, two, or three additional noise variables. All initial guess conversion rates for our approach were sampled around 1 with a perturbation range of [0.2], including the noise parameters, which have to be calibrated towards zero.

As shown in Fig. 4(a), parameter improvement remained largely stable for the upstream parameter *c*_*A*_, but decreased for later parameters in the chain, most clearly for *c*_*E*_. This indicates that additional noise variables mainly reduce the identifiability of downstream parameters, consistent with signal being diverted from the main chain before reaching later states.

**Figure 4:**
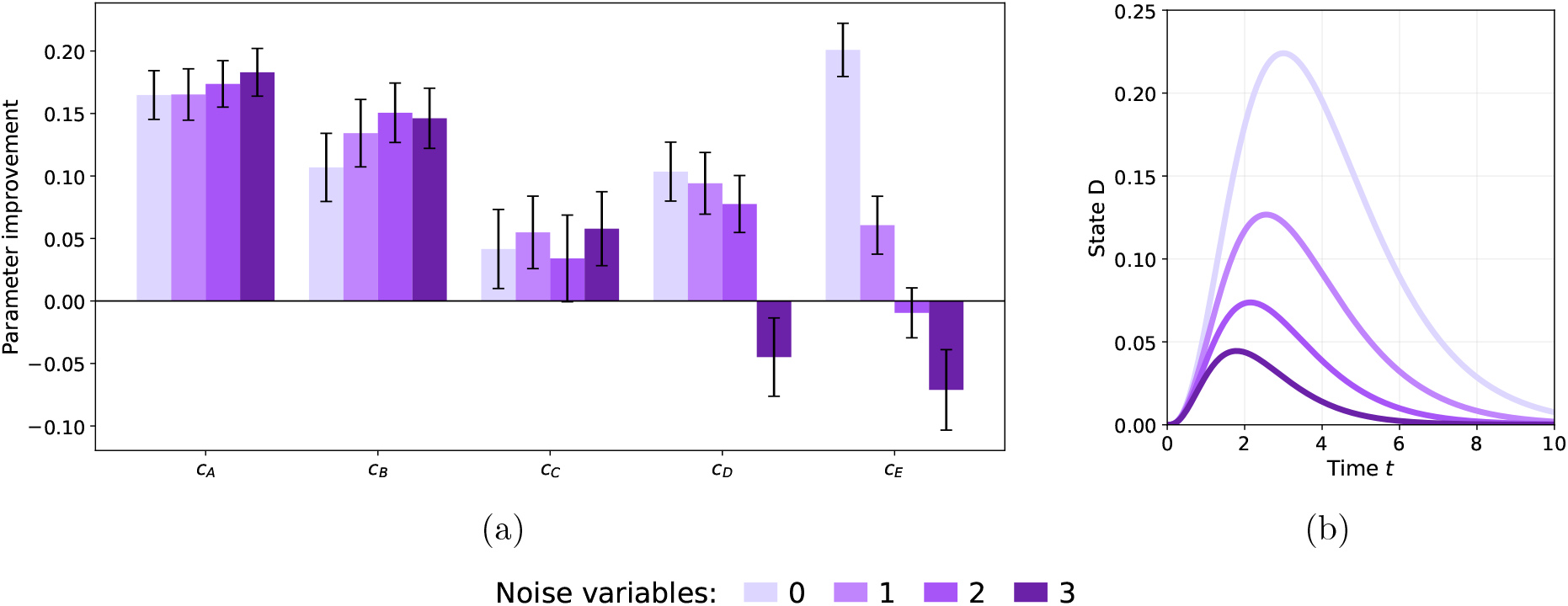
Effect of noise variables on ABCDE calibration. **(a)** Parameter improvement for the core ABCDE parameters *c*_*A*_–*c*_*E*_ after calibration with 0, 1, 2, or 3 additional noise variables. Positive values indicate that the calibrated parameter is closer to the true value than the initial parameter. **(b)** Simulated trajectory of state *D* for the base ABCDE model and extended models with 1, 2, or 3 additional noise variables.

The effect of the noise variables on the ODE dynamics is illustrated in Fig. 4(b). With each added noise variable, part of the signal is diverted away from the main chain, which changes the trajectory of downstream states and makes their parameters harder to recover.

Despite this reduced parameter identifiability, the trajectory RMSE before and after calibration showed only a small impact of the added noise variables (Supplementary Fig. S1), indicating that lack of parameter identifiability is more of an issue of the amount of information contained in the noisy model rather than performance of our specific calibration. Thus, the calibration workflow remains able to improve trajectory agreement in the presence of noise variables, while parameter recovery can become more sensitive to increasing model complexity in some settings.

### Latent Space Separates Distinct Dynamics

A useful latent space for calibration should preserve dynamically relevant differences while reducing the dimensionality of the observed trajectories to enable reasoning about patient subgroups. We therefore examined whether synthetic patient trajectories generated from different ODE parameter regimes become easily distinguishable after encoding.

We simulated this by training the autoencoder on trajectories generated with the parameters *c*_*A*_ = *c*_*B*_ = *c*_*C*_ = *c*_*D*_ = *c*_*E*_ = 1 and encoding trajectories from this parameter set and from a second parameter set with identical values except for *c*_*E*_ = 3, using the same initial conditions and observation times for both groups.

As shown in Fig. 5, the two parameter regimes form visually separable trajectory patterns in latent space. This indicates that a useful autoencoder representation is not limited to its training regime, but still captures dynamic differences in patient trajectories beyond the training set.

**Figure 5:**
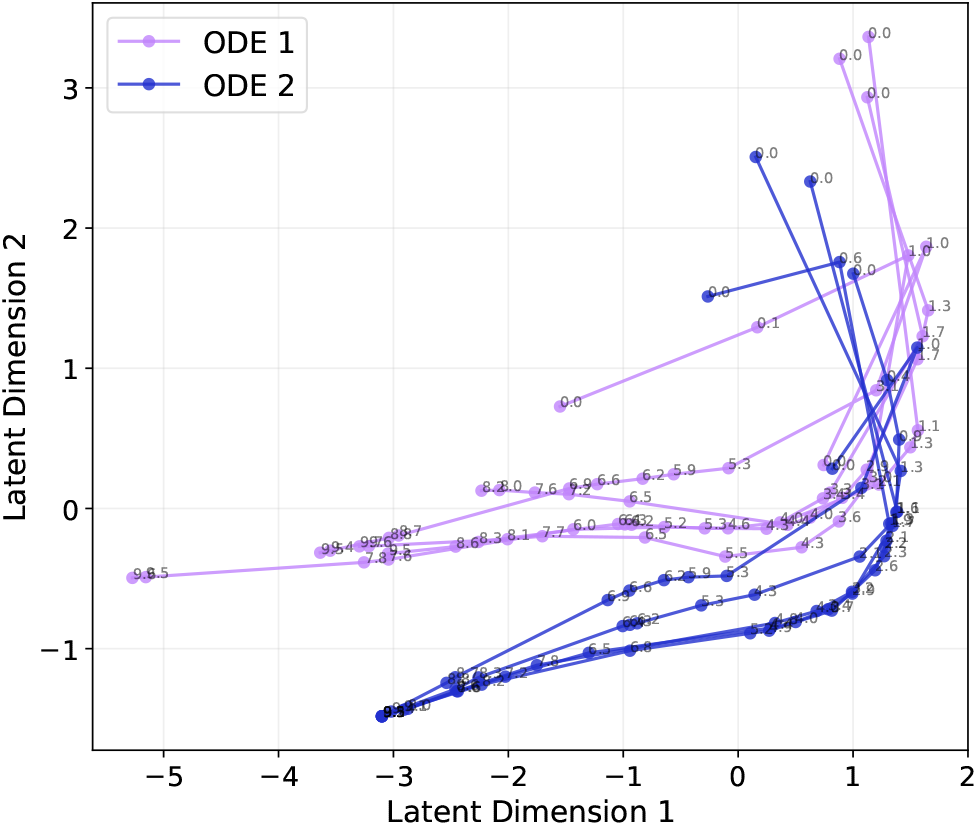
Latent-space representation of simulated ABCDE patient trajectories from two parameter regimes. Both groups were generated with identical initial conditions and observation times, but differed in the parameter *c*_*E*_ (*c*_*E*_ = 1 versus *c*_*E*_ = 3). Trajectories from the two regimes form visually separable patterns in the two-dimensional latent space.

### Calibration of Clinical Epidermolysis Bullosa Data

In cooperation with clinical experts and modelers, we developed a phenomenological model structure for Epidermolysis Bullosa, shown in Fig. 2(b). The corresponding ODE system is given in Eq. (2)) in Methods. The model describes worsening inflammation as the driver for a clinical worsening patient state, as food intake becomes painful, inflammatory processes worsen anaemia, and additional negative consequences deteriorate general patient health. When the general health state is high, patients exhibit stronger resistance to inflammation, but mathematically, the model guarantees deterioration of the health status by another, independently acting mode of increase for inflammation. The model consist of the four variables inflammation *I*, nutrition *N*, anemia state *A*, and general health state *H*, which were mapped to the clinical proxy variables Creactive protein, albumin, hemoglobin, and BMI relative to age- and sex-matched WHO reference values, respectively.

Having defined the model structure and its mapping to observable clinical variables, we applied the calibration workflow to longitudinal data from 19 Epidermolysis Bullosa patients. For this simulation, we started with an initial set of ODE parameters that match our expectations about the time-scales of the relaxation towards the current setpoint (Tab. 1). In order to curb the impact of outliers on standardization, we smoothed data with a moving average prior to calibration.

**Table 1:**
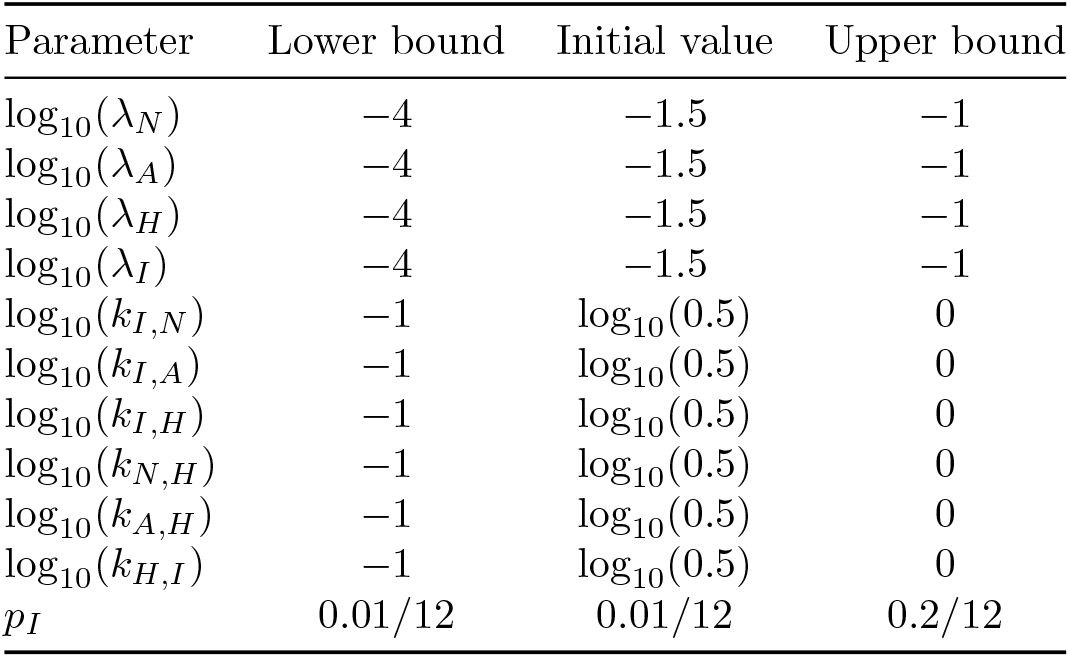
Initial ODE parameter values and calibration bounds for the clinical Epidermolysis Bullosa model. Values are shown on the log_10_ scale used during calibration, except for *p*_*I*_.

| Parameter | Lower bound | Initial value | Upper bound |
| --- | --- | --- | --- |
| $\log_{10}(\lambda_N)$ | -4 | -1.5 | -1 |
| $\log_{10}(\lambda_A)$ | -4 | -1.5 | -1 |
| $\log_{10}(\lambda_H)$ | -4 | -1.5 | -1 |
| $\log_{10}(\lambda_I)$ | -4 | -1.5 | -1 |
| $\log_{10}(k_{I,N})$ | -1 | $\log_{10}(0.5)$ | 0 |
| $\log_{10}(k_{I,A})$ | -1 | $\log_{10}(0.5)$ | 0 |
| $\log_{10}(k_{I,H})$ | -1 | $\log_{10}(0.5)$ | 0 |
| $\log_{10}(k_{N,H})$ | -1 | $\log_{10}(0.5)$ | 0 |
| $\log_{10}(k_{A,H})$ | -1 | $\log_{10}(0.5)$ | 0 |
| $\log_{10}(k_{H,I})$ | -1 | $\log_{10}(0.5)$ | 0 |
| $p_I$ | 0.01/12 | 0.01/12 | 0.2/12 |

Fig. 6 shows that our calibrated trajectories fit the observed data and considerably improve upon the trajectories simulated from initial guess parameters. Across patient data, inter-patient heterogeneity dominates over the slow-moving intra-patient changes, highlighting the necessity of standardization prior to calibration. Although our model fits the data reasonably, this large variability between patients suggests that the used ODE model should incorporate patient baseline differences even more explicitly to get more accurate patient-level predictions.

**Figure 6:**
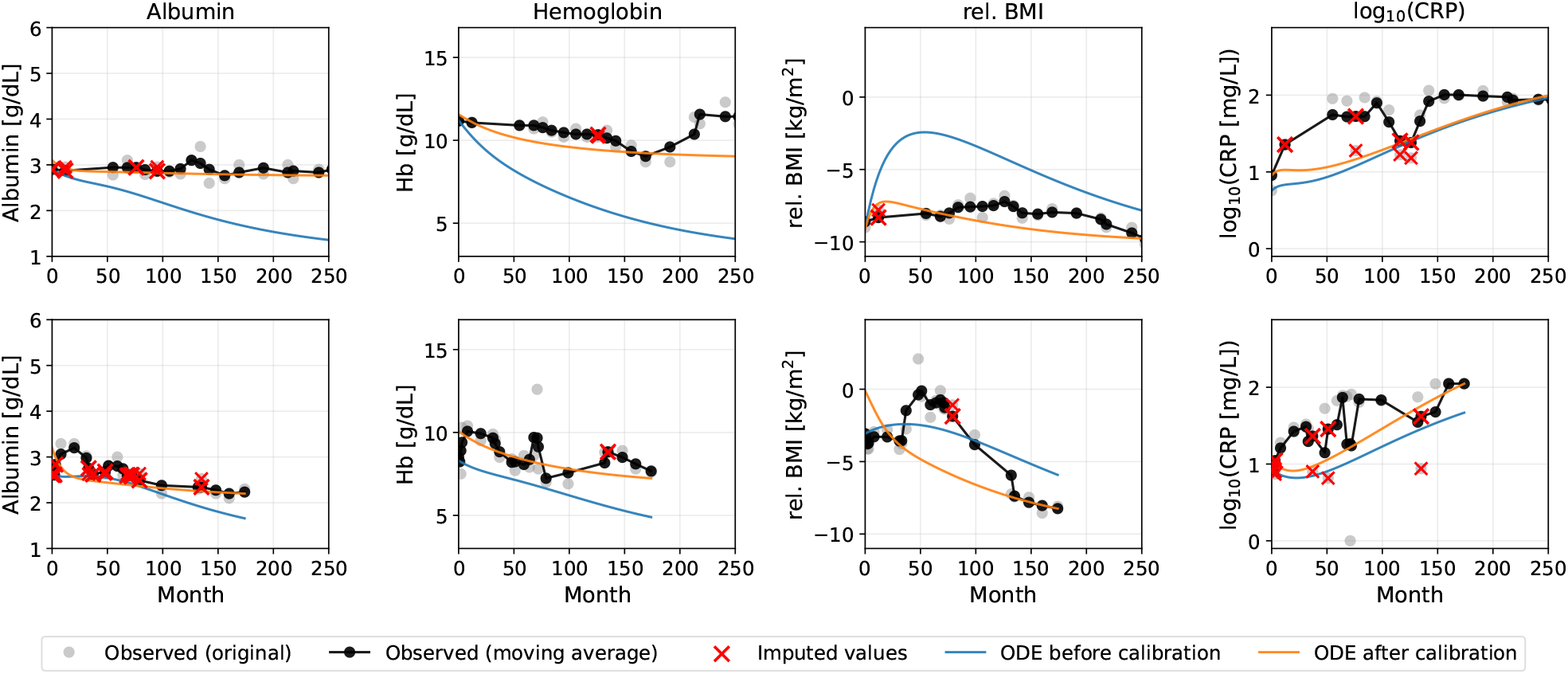
Calibration results on clinical data from Epidermolysis Bullosa patients. Compared to the initial parameter guess for the ODE, our calibrated parameters describe the real data more accurately. Data has been smoothed with a moving average with window size 3 before applying the calibration procedure. Ranges for each clinical variable mirror the observed range across the whole patient set.

## Discussion

Data scarcity for patients suffering from rare diseases does not only limit our qualitative understanding of disease processes, but also hinders quantitative estimation of disease courses or effects of possible treatments. The lack of qualitative understanding produces inexact models that may contain uninformative clinical predictors, while the lack of quantitative data complicates the calibration of the inferred models. These gaps in our ability to research such diseases unfortunately lead to real consequences in terms of patient outcomes.

We have shown in this study that our model calibration in latent space addresses these qualitative and quantitative issues of rare disease modeling simultaneously. By consulting clinical experts for the disease in question, clinical variables informative for the disease at hand can be translated into a mathematical model that phenomenologically links these variables. While such a model might be partially incorrect, we have shown that our latent space approach stays robust under model misspecification, alleviating problems caused by our limited understanding of the disease.

The prior knowledge contained in the model structure allows for high-scale simulation of synthetic patient data. This synthetic data enabled our autoen-coder to feasibly learn the correlations between clinical variables that occur over the course of the disease. Additionally, we also utilized this prior knowledge to impute missing data with a higher degree of accuracy than model-agnostic approaches could do.

Learning the latent representations leaned heavily on per patient standardization of the clinical variables instead of global standardization approaches. Clinical routine data as well as clinical laboratory values were reported to be interpreted with respect to the individual patient references [22] and biological systems more generally respond to relative changes rather than absolute changes [23]. In our approach, patient standardization assumes multiple functions: the baseline shift mainly removes the strong inter-patient heterogeneity at baseline, but the rescaling further reduces the information content from the ODE outputs to the dynamically relevant information.

Patient standardization and dimensionality reduction encourage parsimony, which can be linked to generalizability of the learned model [20, 24]. As our learned representation needs to generalize past the initial guess ODE, learning the standardized ODE solution that contains the relevant elements shared between parameter sets is more likely to generalize to other parameter settings. We suspect that generalization is further enhanced by learning time information not through explicit encoding, but implicitly through the evolving relative state of variables to each other. However, such a claim would need to be substantiated by further research.

Other approaches [20, 21] have also used autoencoders in conjunction with ODEs to generate useful latent representation. The conceptual difference to our approach is that the ODE is defined in the latent space rather than outside of it, with one argument being that “autoencoders generally do not take dynamics into account and, thus, may not provide the right basis for parsimonious dynamical models” [20]. Based on our results we provide concrete evidence to the contrary: our encoded trajectories respect the order of time points well even when this was not an explicit training objective.

Our learned representation has distinct benefits based on its construction from an expert-informed ODE based on heterogeneous synthetic patients. The feasibility of our calibration in latent space suggests that the representation is meaningful beyond the parameters that were used for training the ODE. This opens up possibilities for classifying different patient trajectories on a simple visual level by inspecting the learned trajectories in the low-dimensional latent space: if patient trajectories behave differently, this can be detected already in a human-friendly exploratory manner.

Our study also includes a novel phenomenological model for Epidermolysis Bullosa. The model has been developed in cooperation of clinical experts and a modeler and describes inflammation as the primary driver of a clinical worsening of health-related states, while higher levels of patient health act as protection against worsening inflammation. By building the phenomenological model based on the four qualitative states *inflammation, anaemia, nutrition, general health* and quantitatively linking them to known relevant clinical variables [2], we facilitate interpretation of the model structure without limiting our ability to quantitatively analyze the data. We further simplify interpretation by sticking to a consistent dynamical description through state-dependent set points to which variables will slowly converge to, yielding a specifically natural parametrization of the problem.

We also report limitations that we have observed with our approach. For one, model misspecifications can only be fixed when the amount of perturbations compared to the real data-generating model is limited. Specifically, models can also be wrong in the direction of missing important clinical variables: in this sense, we are still limited by data availability just as much as any other approach. Approaching the problem of accurate model specification might soon be supported though agentic knowledge extraction workflows tailored to ODEs [26, 27], although involvement of clinical experts will still provide crucial context during the model search.

Another limitation is observed in the noticeable drop off in imputation performance if the true model parameters diverge too far from the initial guess parameters on which the imputation algorithm was trained. One methodical improvement would be to also draw the dynamic parameters from distributions to better generalize out of sample. While we have generally observed the autoencoder to be much more robust to parameter misspecification, having a poor initial guess ODE will similarly limit performance for this module. One possible remedy to parameter misspecification is an iterative recalibration of autoencoder and imputation algorithm based on the updated ODE every couple of iterations. We might also consider a wholistic approach that optimizes all components together, although it is conceptually not obvious how to orchestrate such an algorithm given the need for updated synthetic data augmentation for our sparse clinical dataset.

Overall, this study reports an approach that calibrates expert-derived ODEs for rare diseases within a latent space, augmenting the scarce patient data with synthetically generated data. Our approach effectively deals with the problem of misspecified ODEs by learning compact, generalizable representations with a synthetically calibrated autoencoder that filters out irrelevant clinical variables by detecting mismatches between the ODE and the available data. This approach was extensively verified on a model with simulated data and further validated on an expert-derived model for Epidermolysis Bullosa with real data from only 19 patients. With already promising results in this study, further extensions to this approach are conceivable.

## Methods

### Synthetic data generation

ODEs describe the change of a state based on the set of state values at any given time with a set of equations and model parameters that define the exact relationship. States might represent the abundance of a certain protein or gene transcript in a cell, or more clinical variables such as weight or hemoglobin values in the blood. Given states *x* and a set of parameters ***θ***, we can describe a set of differential equations as

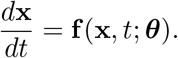

We assume that we can observe values *x*_*real*_ for the states *x* directly, in contrast to the more general setting of using observation functions [28]. However, we allow that a state appears in the equation although never being observed, allowing for a more flexible causal representation based on a given set of clinical variables.

We use this ODE to simulate synthetic data to train an autoencoder and imputation algorithm later. To simulate synthetic data based on a heterogeneous population of patients, we introduce a distribution of their initial conditions. In real data settings, this distribution of initial conditions can be estimated based on the observed data, including a possible correlation matrix between clinical variables. In order to simulate useful synthetic data, we require that the parameters *θ*_0_ are in the approximate vicinity of the data-generating parameters *θ*_*true*_; the extent to which this matters is demonstrated in the results.

The generation of synthetic data thus proceeds as follows. First, we draw a set of initial conditions to represent different synthetic patients. For these, we simulate the ODE to obtain their full time trajectories and then randomly sample a fixed amount of time points from this ODE to obtain 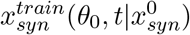; these data are used to train the autoencoder. In instances in which we want to have our synthetic data aligned with the real data as much as possible, we fix the initial conditions to the ones of the real data and sample the full time trajectory at the time points of the corresponding real patients to obtain 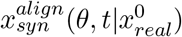; this data is used when comparing real data to synthetic data.

### Autoencoder

To compare the measured data *x*_*real*_ to the corresponding trajectories of the ODE in the latent space, we first learn a map from the observed space into the latent space with an autoencoder based on the synthetic training 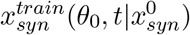 that is sampled as described above.

The autoencoder training is visualized in Fig. 1(a). We first draw synthetic data from the initial ODE with randomly sampled initial conditions and the initial guess for the dynamic parameters *θ*_0_. The clean synthetic patient trajectories are sampled at random time points 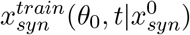, noise is added to them, all clinical variables are patient-standardized 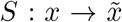 and propagated through the autoencoder 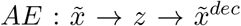 with encoder *Enc*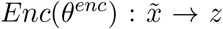 and decoder *Dec*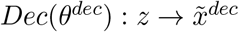for many randomly drawn initial values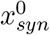:

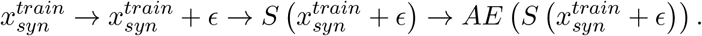

We then optimize the autoencoder parameters by aligning the output of the standardized, noisy synthetic data to the clean standardized ODE solution.

While standardization of variables is a standard technique when using autoencoders, the clinical setting of inter-patient variabilities that are considerably higher than intra-patient dynamics requires a patient-wise standardization approach. This is necessary because the high inter-patient variability prevents the autoencoder to learn the correlations between clinical variables, because a high value of a variable for a patient may be either due to being at a late stage of the disease process or due to a high baseline value for the patient. Thus, for each patient, we standardize each clinical variable with the map 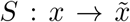 to the fixed range of [-1,1].

Our trained autoencoder achieves two objectives. First, by constraining the latent space to a lower dimensionality, the autoencoder is forced to keep only informative structure contained in the relationships between clinical variables, filtering out noise and uninformative variables. Second, by drawing synthetic patients from many different initial conditions, the patient-standardization forces the autoencoder to learn dynamic correlation between clinical variables more generally compared to encoding absolute values directly, increasing the generalizability across initial conditions and dynamical parameters.

### Calibration

The ODE calibration in latent space aims to find ODE parameters *θ* such that the distance between patient data *x*_*real*_(*t*) and the corresponding synthetic data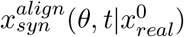 is minimized. The iterative calibration procedure is illustrated in Fig. 1.

The optimization is initialized with the prespecified initial guess parameter values *θ*_0_ that were used to train the autoencoder. For each calibration itera-tion, synthetic trajectories 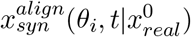 are generated with the current ODE parameters *θ*_*i*_, using the same initial conditions and observation times as the observed patient data *x*_*real*_. Both observed and synthetic trajectories are standardized and passed through the trained encoding network. The calibration loss is defined as the RMSE between the corresponding latent representations

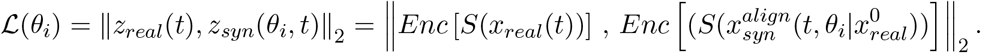

The loss is differentiable with respect to *θ* through the ODE solution and the encoder, allowing the ODE parameters to be updated through gradient-based optimization. This procedure is repeated for a fixed number of calibration epochs, resulting in a calibrated parameter set *θ*_*cal*_ with minimal loss.

### Imputation

Observational data from clinical settings are usually sparse and contain missing values. Since the autoencoder requires complete input vectors, missing values must be imputed before latent-space calibration. To handle this, we employ an imputation approach that is built on a-priori knowledge from the initial ODE system.

We train a mask-aware multilayer perceptron on synthetic data 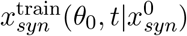 generated from the ODE model. During training, complete synthetic trajectories are masked assuming missing completely at random (MCAR) mechanism. The masked trajectories are then transformed using the same patient-wise standardization described for the autoencoder training. The imputer receives both the partially observed clinical-variable values and the corresponding missingness mask as input and is optimized to reconstruct the corresponding ODE solution from the masked entries.

### ABCDE Model

The base ABCDE model shown in Fig. 2(a) consists of five states *A*–*E* connected in a unidirectional chain:

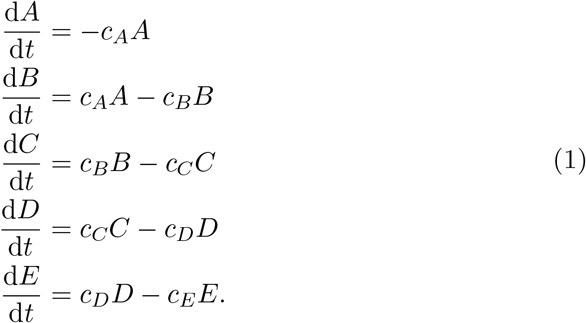

Each parameter *c*_*A*_, …, *c*_*E*_ controls the outflux from the corresponding state and, except for *E*, the influx into the subsequent state.

We adapted this model for our analysis of noise variables by adding alternative outfluxes into the model, but setting the corresponding conversion parameters to zero. An example of how the noise state *B*^′^ interacts with the equation for *B* would be:

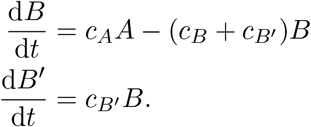

Additional noise states were introduced analogously for later states in the chain.

### EB Model

EB is a rare inherited skin fragility disorder characterized by blistering, erosions, and chronic wounds after minor mechanical trauma. In severe forms, EB can be accompanied by systemic complications including chronic inflammation, impaired nutritional status, anaemia, and severe weight loss [29]. While a fully mechanistic patient-level disease model can not feasibly be inferred from available data, availability of long-term observational data enabled the construction of a phenomenological model. This model represents patient nutrition *N* (albumin), anaemia resistance (hemoglobin), inflammation *I* (C-reactive protein) and general health state (BMI relative to healthy population) through the following equations:

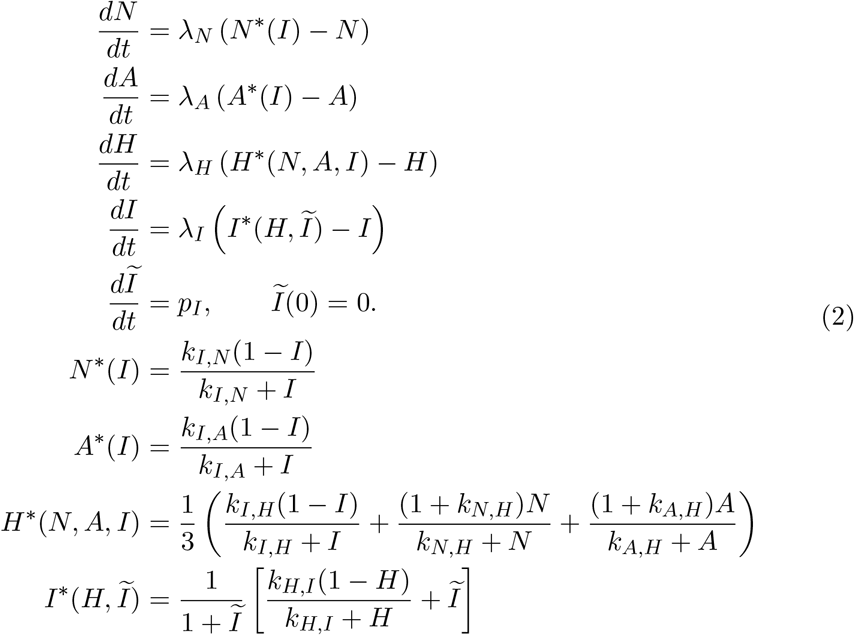

To simplify parametrization, all dynamic variables were standardized to a range of [0,1] across the whole patient population. The model relaxes each variable toward a set point determined by the other variables with rate *λ*_*N*_, *λ*_*A*_, *λ*_*H*_, or *λ*_*I*_. The auxiliary state *Ĩ* represents the unobserved progressive inflammatory drive and is not used as an observed clinical variable. The initial guess parameters for this model are specified in Tab. 1.

## Acknowledgements

Funded by the Deutsche Forschungsgemeinschaft (DFG, German Research Foundation) – Project-ID 499552394 – SFB 1597.

## Data availability

Synthetic data used for method evaluation can be generated using the code provided with this article. The clinical Epidermolysis Bullosa data are not publicly available because they contain patient-level rare-disease data subject to privacy and ethical restrictions. Access to these data may be requested from the corresponding author and is subject to institutional approval and applicable data-use agreements.

## Code availability

The Python implementation of the latent-space ODE calibration workflow is available at https://gitlab.imbi.uni-freiburg.de/timo.rachel/CalibratingODEsInLatentSpace. The repository contains a Jupyter notebook and reusable Python modules for synthetic data generation, autoencoder training, ODE-informed imputation, latent-space ODE calibration, and evaluation. An archived version with a persistent identifier will be provided upon publication.

## Author contributions

HB, MH and HY conceived the initial study idea, with further conceptual development by HY, TR and TL. HY and TR implemented the computational workflow and performed the analyses. JT contributed expertise in ODE modeling. MK, AR-T and CH provided clinical expertise and contributed the clinical data. HY, TR and TL wrote the manuscript. All authors reviewed and approved the manuscript.

