## Supplementary Material for "Synthetic-data augmented calibration for expert-informed rare disease models"

6th May 2026

### Additional Figures

#### Calibration Remains Robust to Noise Variables

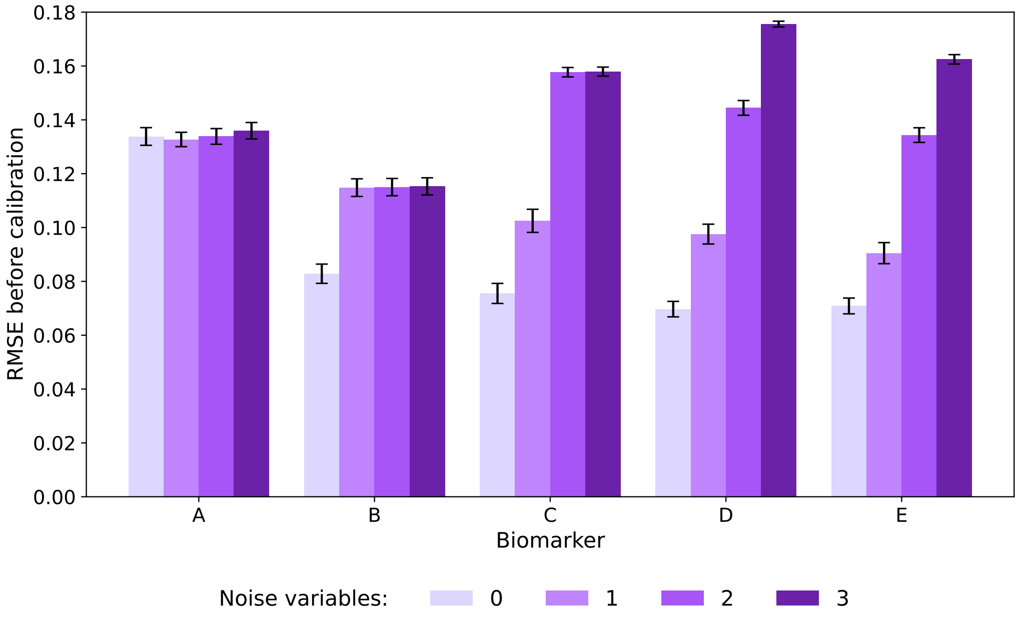


(a) Before calibration

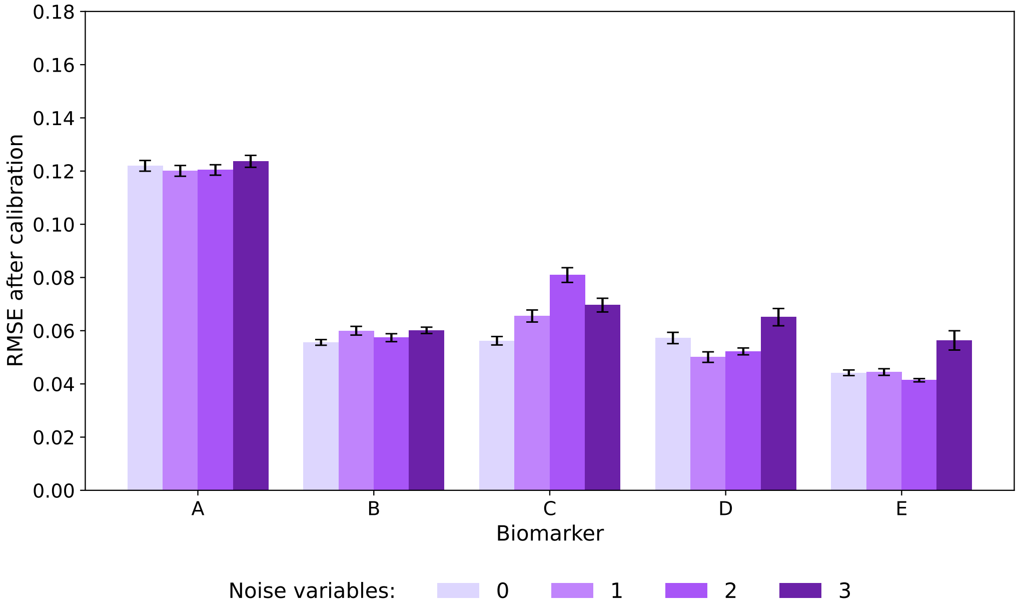


(b) After calibration

Figure S1: **Trajectory RMSE is only weakly affected by additional noise variables.** Biomarker-wise trajectory RMSE for the core ABCDE states $A$–$E$ before calibration **(A)** and after calibration **(B)** in models with 0, 1, 2, or 3 additional noise variables. Bars show the mean across repeated runs and error bars indicate the standard error of the mean. Although additional noise variables reduce downstream parameter identifiability, their effect on trajectory RMSE remains small, and calibration continues to improve agreement between simulated and target trajectories across model complexities.
